# Gamma-burst cortical activity in awake behaving macaques

**DOI:** 10.1101/2023.09.26.559594

**Authors:** Benjamin T. Acland, Ben Julian A. Palanca, Janine Bijsterbosch, Lawrence H. Snyder

## Abstract

Electrophysiological recordings during ketamine anesthesia have revealed a slow alternating pattern of high- and low-frequency activity (a “gamma-burst” pattern) that develops with the onset of general anesthesia. We examine the role of NMDA receptor antagonism in generating the gamma-burst pattern and the link between gamma-bursts and dissociative anesthesia. We compare the effects of ketamine with those of the highly selective NMDA receptor antagonist CGS 19755 on multi-site intracranial electrophysiology and behavior in rhesus macaques. Remarkably, we find that animals given a moderate dose of CGS 19755 are able to perform a difficult memory task, while at the same time showing electrophysiological activity similar to ketamine anesthesia, with one key difference: a lack of delta-band LFP modulation. This difference demonstrates that ketamine’s ability to drive strong delta-band oscillations relies on additional mechanisms beyond NMDA receptor antagonism alone, and points to a key role for the activity underlying delta-band oscillations in causing anesthesia.

## Introduction

Research into ketamine’s influence on neural activity and behavior dates back to the 1960s. A derivative of phencyclidine, ketamine was developed with hope that it could achieve similar levels of anesthesia without the undesirable side-effects of its parent drug. In higher doses, ketamine induces a state termed “dissociative anesthesia,” marked by a distinctive electrophysiological signature: while most general anesthetics induce a progressive dominance of lower frequency EEG activity through theta (4-8 Hz) and then delta (0.5-4 Hz) frequency bands, ketamine anesthesia is associated with elevated theta-band power and beta activity 1 and a “gamma-burst” pattern of slowly alternating low- and high-frequency activity that appears at the onset of behaviorally-defined unconsciousness ^2,3^. These distinct patterns of ketamine anesthesia relative to GABA-ergic anesthetics are thought to arise from distinct molecular targets of ketamine, which has only minor agonism of GABA receptors. Many behavioral and electrophysiological effects of phencyclidine and ketamine are also elicited through non-competitive NMDA receptor antagonist MK-801 and the competitive NMDA receptor antagonist CGS 19755 (cis-4-phosphonomethyl-2-piperidine-carboxylic acid) ^4^. This raises the possibility, highlighted by Akeju et al. (2016), that NMDA antagonism is the main driver of gamma-burst activity during ketamine general anesthesia^3^. At the same time, there is a growing consensus that ketamine’s dissociative and anesthetic effects are mediated by targets besides NMDA receptors ^5,6^. Thus, it may be that NMDA antagonism alone is sufficient to drive gamma-burst activity strongly associated with ketamine-induced unconsciousness, but that this activity is not itself a reliable indicator of unconsciousness.

In this study, we report gamma-burst activity caused by anesthetic doses of ketamine anesthesia and by *sub-anesthetic* doses of the competitive and selective NMDA antagonist CGS 19755. This finding establishes NMDA antagonism as a driver of gamma-burst activity that is not sufficient on its own to induce general anesthesia.

## Results

We collected multi-channel extracellular voltage recordings from rhesus macaques sitting head-fixed in a dark room before and after intramuscular injections of either ketamine or the NMDA antagonist CGS 19755. Before each recording, six to eight electrodes were lowered into separate sites in cingulate and parietal cortex in both hemispheres (3-4 in each hemisphere). In some sessions, recordings were also obtained from temporal, occipital and frontal cortex. Data were recorded from all electrodes simultaneously without interruption. Each recording session consisted of a one-to two-hour pre-injection period, a single intramuscular injection, and a two to five-hour post-injection period. Ketamine doses were either 3 mg/kg (sub-anesthetic) or 10 mg/kg (anesthetic), and CGS 19755 doses were either 3, 6 or 12 mg/kg (sub-anesthetic). We refer to 3 mg/kg as a sub-anesthetic dose because, although animals become clearly sedated and will not initiate tasks for reward (see below and **Fig. 4**), they remain able to move freely and attend to events in the room. During recordings we monitored high- (500-4000 Hz) and low-pass (0.1-1.0 Hz) filtered local field potential (LFP) signals on all channels using a visual display resembling the top two rows of **Figs. 1A and C**.

**Figure 1.**
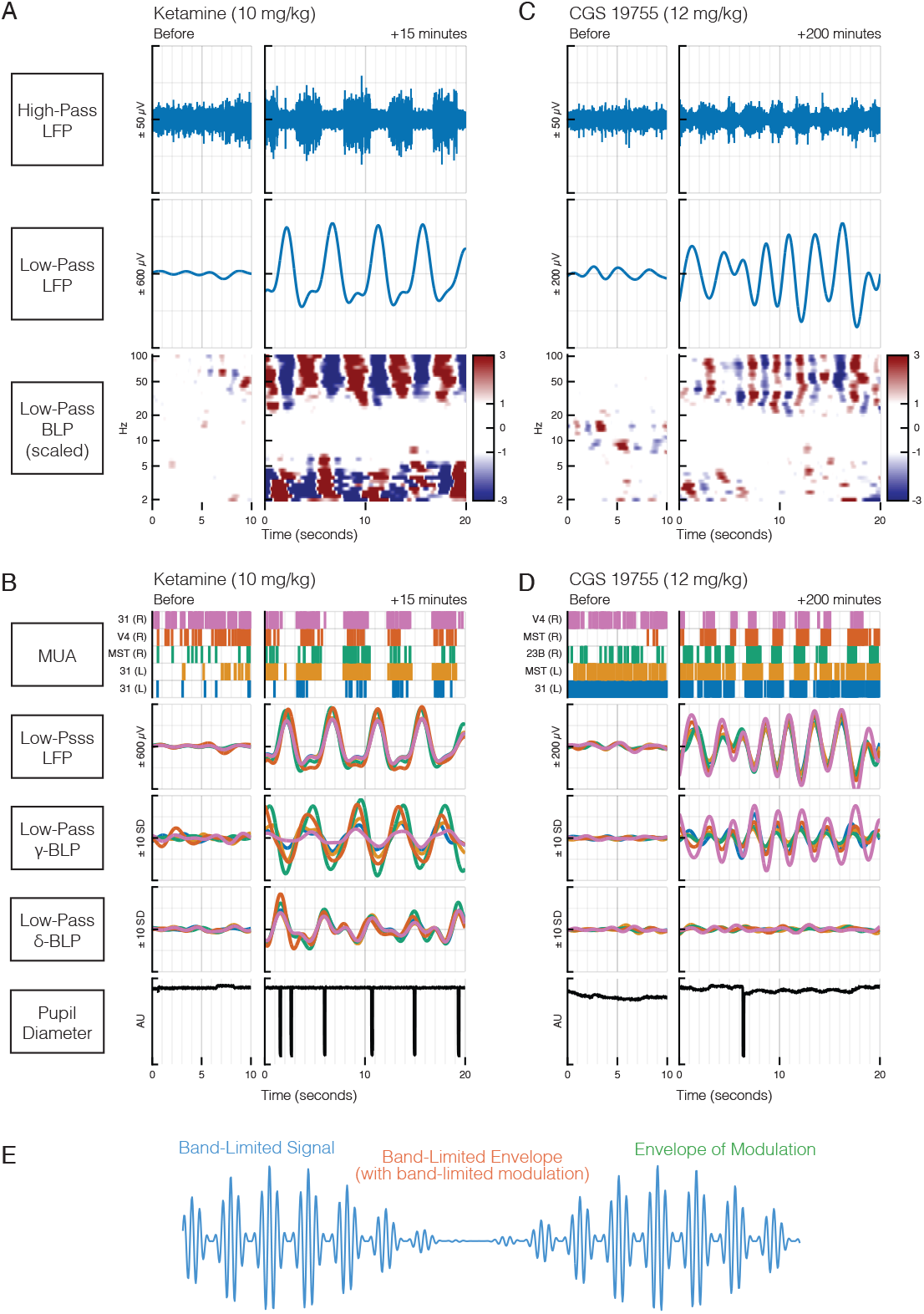
The non-specific NMDA antagonist ketamine and the specific NMDA antagonist CGS 19755 have similar electrophysiological effects in the cortex, driving synchronized oscillations with a period of several seconds. **A**. Ketamine evokes alternating periods of 2.5 s of fast spiking followed by 2.5 s of no spikes (right upper panel, extracellular voltage recording (LFP) filtered at 500-4000 Hz). No such periodicity occurs prior to the ketamine injection (left upper panel). Vertical gray lines are 1 s apart. Low-pass LFP amplitude (right middle panel, 0.1 - 1 Hz) shows a synchronized modulation, low during each burst of spikes and then rising abruptly to peak at the start of each quiet period. Synchronized oscillations are also seen in the low-pass (0.1 - 1 Hz) band-limited power of the LFP, in particular at 40-100 Hz and, in counter-phase, from 2-5 Hz (time-frequency spectrogram, bottom panel; see panel E for details). Intermediate frequencies are not modulated. Values are Z-scored, that is, deviations from the average power in the baseline period are divided by the standard deviation of power in the baseline — see scale bar at lower right. Data for all panels in A are from a sharp micro-electrode in cingulate cortex in the left hemisphere, recorded 10 minutes prior to a 10 mg/kg IM ketamine injection (left) and 15 minutes after injection (right). **B**. Oscillations were not only synchronized across different electrophysiological measures (panel A), but also across locations in cortex. From top to second-from-bottom, multi-unit activity, low-pass LFP amplitude (0.1 - 1 Hz), low-pass gamma (24-60 Hz) LFP band-limited power (see panel E) and low-pass delta (1-4 Hz) LFP band-limited power are not only synchronized with one another, but also across multiple cortical recording sites (area 31 [cingulate cortex] in the right hemisphere, visual area V4 on the R, visual area MST on the right, and two different area 31 sites on the left). See text for additional details. Blinks (pronounced negative excursions in bottom panel) were synchronized with the end of the spike bursts. Recording times identical to panel A. Data in A are from the orange electrode. **C**. CGS 19755 evokes alternating periods of 1 sec of fast spiking followed by 1 s of no spikes (right upper panel). Low-pass LFP amplitude (right middle panel) shows a synchronized modulation, low during each burst of spikes and then rising abruptly to peak at the start of each quiet period. Synchronized oscillations are also seen in the low-pass (0.1 - 1 Hz) band-limited power of the LFP, in particular from 30 to 100 Hz (time-frequency spectrogram, bottom panel; see also panel E). Data for all panels in C are from a sharp micro-electrode in visual area MST in the left hemisphere, recorded 10 minutes prior to a 12 mg/kg IM CGS 19755 injection (left) and 15 minutes after injection (right). Format otherwise similar to panel A. **D**. Oscillations were not only synchronized across different electrophysiological measures (panel C), but also across locations in cortex. Recording sites are indicated at top left. Format is otherwise identical to panel B. **E**. The local field potential (LFP) can be processed in different ways. A *band-limited signal* (blue) is extracted by band-pass filtering the raw LFP amplitude, using both hardware filters (e.g., high-cut anti-aliasing and low-cut filters that limit the dynamic range of the signal) and software (digital) filters. For example, we extract the low-pass LFP by band-limiting the LFP amplitude to 0.1 to 1 Hz. Higher frequencies are also commonly extracted in this way; for example, the gamma band-limited amplitude could be obtained by filtering the LFP from 30 to 100 Hz. The *band-limited envelope* (red) is obtained by measuring the amplitude of each successive cycle of this “carrier” frequency. Squaring this signal produces band-limited power (BLP). In our example, this would be gamma BLP. One can apply this same process a second time, this time to the BLP, to produce an *envelope of modulation* (green) of the BLP. (For simplicity, we show the envelope of modulation of the amplitude, not the power.) The envelope of modulation can contain frequencies from 0 to roughly half the carrier signal. This envelope can be filtered a second time to further limit its frequency content. To compute the low-pass gamma BLP, for example, we filter the gamma BLP from 0.1 to 1 Hz.

Anesthetic doses of ketamine in humans are known to drive a “gamma-burst” pattern of EEG activity — relatively high-power gamma-band oscillations interrupted by slow-delta (0.1-4 Hz) oscillations, repeating at a frequency below 1 Hz ^3^. **Fig. 1A** shows such a pattern evoked by an intramuscular injection of 10 mg/kg ketamine. Multiunit spiking activity cycled between ∼2 s of high rates of firing and ∼2 s of silence. LFP showed similar periodicity, with different effects at different frequencies. At low frequencies (0.1-1 Hz) the LFP voltage (see below for *power*) was low during the high-spike-rate periods and high during the low-spike-rate periods. Peak voltage occurred near the middle of the low-spike-rate period. At higher frequencies, the amplitude of LFP oscillations was synchronized with changes in spiking activity. To capture this modulation we looked at narrow (band-passed) frequency ranges, first computing the envelope of the band-passed signal and then filtering that envelope to extract only power between 0.1-0.6 Hz, that is, the power associated with the gamma-burst pattern (**Fig 1E**). This analysis of filtered band-limited power (BLP) showed that high frequencies of LFP (30-100 Hz — gamma-band) were modulated in phase with spiking activity, low frequencies (2-5 Hz — delta- and low theta-band) were modulated in counter-phase with spiking activity, and intermediate frequencies (5-20 Hz — high theta-, alpha- and beta-band) were unmodulated (**Fig. 1A**, spectrogram).

Not only was this gamma-burst pattern present at all locations tested, but the bursts themselves were synchronized across simultaneously recorded channels sampling across cerebral cortex. An example of this is shown in **Fig. 1B**. Five recording sites, in left posterior cingulate, right posterior cingulate, right occipital, and right temporal cortex, showed synchronized modulation of spiking rate, low-pass LFP, gamma BLP, and delta BLP. Low-pass (0.1-0.6 Hz) LFP is particularly well-aligned across channels, with phase offsets of less than ±4 deg (< ±50 ms). Spike rate shows some cycle-by-cycle variation but when averaged across cycles, phase offsets in cumulative spike histograms are minimal (data not shown). Gamma BLP shows phase offsets of up to ±20 deg. Delta BLP is well-synchronized across channels, though the activity patten itself is more complex than what is seen in the other modalities, with a trough in power coincident with the midpoint of the burst in spike activity and peaks at the start and end of the burst. Blinks were synchronized with the end of spike bursts.

CGS 19755, a more selective NMDA antagonist, also produced gamma-burst activity, with many similarities and some differences from the pattern produced by ketamine. A 12 mg/kg CGS 19755 dose produced a gamma-burst pattern in spikes and LFP with high-spike-rate periods lasting ∼1 s and low-spike-rate periods lasting ∼1 s (**Fig. 1C**), about half the period of the gamma-burst activity seen following a 10 mg/kg ketamine dose. Low-pass LFP voltage was synchronized with spiking activity similar to what was seen under ketamine, with a slightly earlier peak voltage coinciding with burst offset. LFP BLP was modulated in phase with spike rate for beta and gamma bands (20-100 Hz, strongest from 30-70 Hz), but modulation was poor or absent for lower bands including delta.

Like ketamine, CGS 19755 induced this gamma-burst pattern at all locations tested, with some key differences. An example shows the patterns in posterior cingulate, temporal, and occipital cortex (**Fig. 1D**). Spiking was aligned across all five channels in which unit activity was recorded. Three channels were clearly synchronized. Two continued spiking during the low-spikes phase of some but not all intervals. Low-pass LFP showed clear oscillations, all at the same frequency, with phase offsets of ±4 deg. Gamma BLP showed synchronization with offsets < ±20 deg. Critically, delta BLP showed little modulation or synchronization in stark contrast to gamma-burst activity under ketamine. Similar patterns were seen at lower doses, with slightly more cross-channel phase offset in low-pass LFP and no clear synchronization of delta BLP (**Sup. Figs. 2-3**). Blinks did not synchronize with neural activity at any dose of CGS 19755 administered in this study.

While the gamma-burst patterns evoked by ketamine and CGS 19755 had many similarities, the time courses of the associated low-frequency signal modulations were quite different between drugs. **Figure 2 A** and **B** show the injection-aligned average of low-frequency (0.1 to 1 Hz) modulation power over time for spikes, LFP voltage, and LFP BLP. Modulation power in spikes and low-pass (0.1 to 1 Hz) LFP was quantified by measuring power in the 0.1 to 1 Hz range. Modulation power in BLP was quantified by measuring power in the low-pass envelope (0.1 to 1 Hz) of delta and gamma BLP. Data for each signal were normalized using their pre-injection modulation intensity (see Methods), then averaged over recording sites (26 sites from 6 sessions for 10 mg/kg ketamine; 17 sites from 4 sessions for 12 mg/kg CGS 19755).

**Figure 2.**
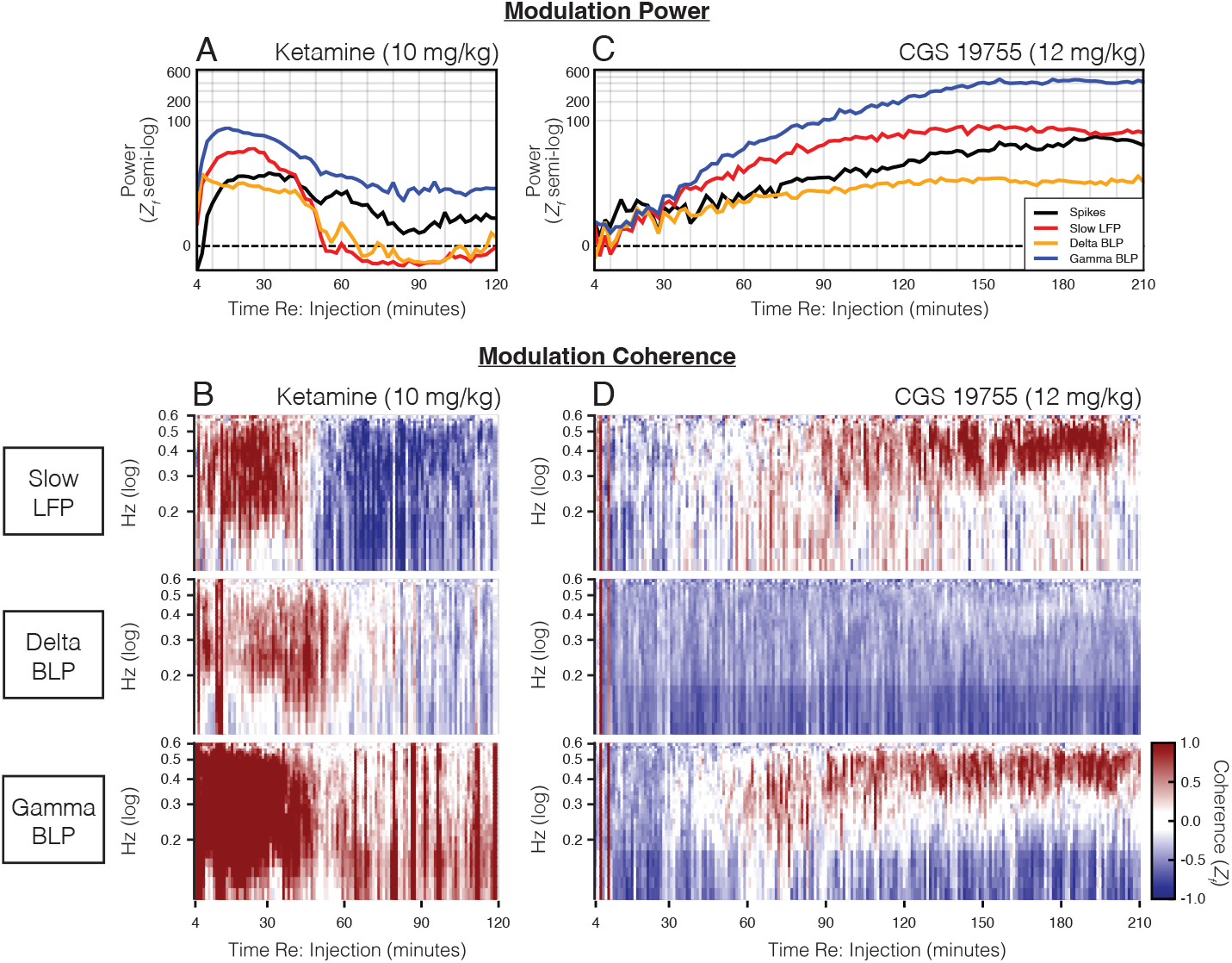
Ketamine and CGS 19755 induce a similar “gamma-burst” pattern including slow (> 1 s period) oscillations in various electrophysiological measures, but the time course of modulation is markedly different for the two drugs. **A**. Slow oscillations are quantified as signal power from 0.1 to 1 Hz. Ketamine increases power in spike rate (black), low frequency LFP amplitude (“slow LFP”, red), low frequency band-limited LFP power (delta BLP: 2-5 Hz, yellow), and high frequency band-limited power (gamma BLP: 30-100 Hz, blue), each measured between 0.1 and 1 Hz (see Fig. 1E), by a factor of 10 to 100. This increase in low frequency power, which indexes the slow oscillations, begins almost instantaneously and peaks within 30 minutes. Oscillations in delta and gamma BLP are suppressed below baseline levels after 1 hour, while spike and gamma BLP oscillations drop from the early peak but remain above baseline for at least 2 hours. **B**. Slow oscillations are synchronized across recording sites in all of these signals. Synchrony is quantified as changes in coherence between pairs of electrodes, relative to baseline coherence. Coherence is plotted from 0.1 to 0.6 Hz as a function of time. Peak coherence occurs from 0.2-0.5 Hz. The time course of coherence follows the time course of power: when power is elevated, coherence is also elevated, and when power falls below baseline (>1 hour, slow LFP and delta BLP), coherence also falls below baseline. Changes in coherence are scaled to baseline coherence SD (scale bar, bottom right of panel D). **C**. Like ketamine, CGS 19755 increases the power of multiple electrophysiological measures between 0.1 and 1 Hz, corresponding to slow oscillations in these signals. Unlike ketamine, the effect starts small and grows steadily over the next 2-3 hours. Slow oscillation power plateaus ∼2.5 hours post-injection, and the plateau is maintained for at least another hour. Format similar to A. Note that despite the difference in time course, the order of the traces is the same for the two drugs (compare panels A and C). **D**. Slow oscillations show increasing synchronization across recording sites as time elapses, with peak coherence at 0.4-0.5 Hz. Format similar to B. In panels A-D, plots start at 4 min rather than 0 to avoid alerting effects and movement artifacts

Ketamine effects appeared within minutes of injection, with rapid increases in modulation power for spikes, low-pass LFP and BLP. Delta BLP modulation power reached its peak value ∼5 minutes after injection, followed by gamma BLP and low-pass LFP at ∼15 minutes, and finally spikes at ∼20 minutes. Low-pass LFP and delta BLP modulation power decreased gradually until ∼50 minutes, beyond which point they fell off rapidly and were suppressed below baseline for another ∼60 minutes. Gamma BLP and spike modulation power remained near their peaks until ∼40 minutes after injection and then decreased gradually, remaining elevated until at least 120 minutes after injection (when recordings stopped).

Gamma-bursts were synchronized across all sites (**Fig. 2B**). To quantify the synchrony of signal modulations, the coherence of low-pass LFP and the low-pass (0.1-1 Hz) envelope of delta and gamma BLP was computed between every pair of simultaneously recorded sites. Data for each signal pair were normalized using their pre-injection modulation intensity, then averaged across all pairs (see Methods). The time course of coherence matched the time course of modulation power (**Fig. 2A**). Coherence in the low-pass LFP and the low-pass envelopes of delta and gamma BLP rose early and remained strong for 40-60 minutes. Low-pass LFP and delta BLP coherence then fell off rapidly, with low-pass LFP coherence dropping below pre-injection levels and remaining low until recording stopped ∼120 minutes after injection. Though gamma BLP coherence also fell off after 40-60 minutes, it remained moderately elevated for at least 2 hours.

CGS 19755 had similar effects on modulation power, but with a very different time course (**Fig. 2C**). In contrast to ketamine’s rapid onset, the effects of CGS 19755 began to appear only ∼20 minutes after injection, increased until 130-150 minutes, and persisted at high levels for at least 220 minutes (when recordings were stopped). This difference in time courses likely reflects different rates of crossing the blood-brain barrier, metabolism and excretion ^7-11^. The coherence of signal modulations caused by CGS 19755 followed the time course of modulation power for low-pass LFP and gamma BLP, but in contrast to ketamine this was not the case for delta BLP, which instead showed slightly reduced coherence compared to the pre-injection period (**Fig 2D**). For each drug, similar patterns were seen at lower doses (**Sup. Figs. 4-6**).

**Fig 1** shows a difference in the dominant frequency of gamma-burst activity caused by 10 mg/kg ketamine (∼0.25 Hz) and 12 mg/kg CGS 19755 (∼0.4 Hz). This difference may reflect a dependency of the frequency of modulation on the fraction of NMDA receptors bound: while these doses contain similar numbers of molecules (42 versus 54 mmol/kg, respectively), ketamine has substantially higher bioavailability ^7-11^. To test this hypothesis, we compared changes in the dominant frequency of modulation for a ketamine dose of 10 mg/kg and CGS 19755 doses of 3, 6 and 10 mg/kg (**Fig. 3** and **Sup. Figs. 7-9**). We calculated the dominant frequency of modulation over time by applying an autocorrelation-based approach to 5-minute windows of data beginning after injection (see Methods). Data were then averaged across recording sites for each dose. While modulation power for each drug changed over time (**Fig. 2** and **Sup. Figs 4-6**), the dominant frequency of modulation was fairly stable over time. With increasing doses of CGS 19755 the dominant frequency of modulation decreased, from ∼0.55 Hz for 3 mg/kg to ∼0.5 Hz for 6 mg/kg and ∼0.4 Hz for 12 mg/ kg. Signal modulations evoked by a ketamine dose of 10 mg/kg were at ∼0.25 Hz. This decrease in modulation frequency is consistent with an inverse relationship between the fraction of NMDA receptors blocked and the frequency of modulation.

**Figure 3.**
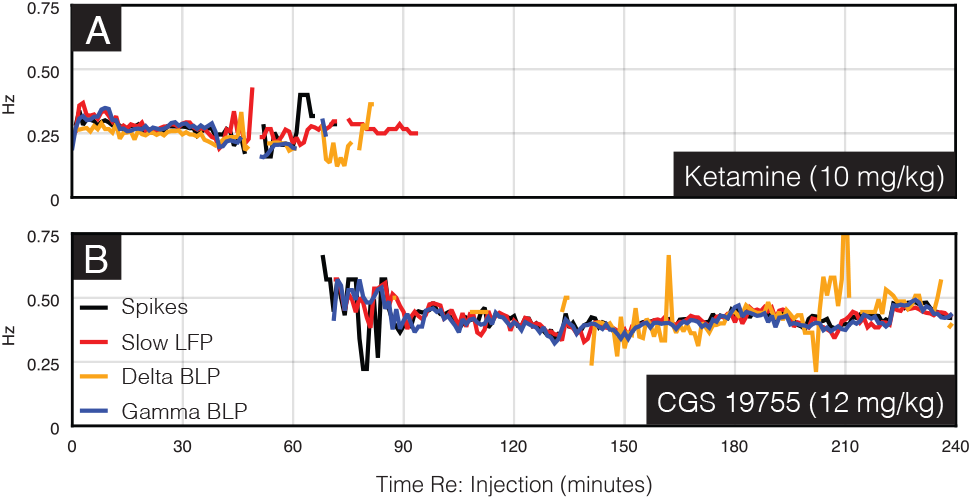
The dominant frequency of slow oscillations in 4 electrophysiological measures (see Fig. 2) are similar but not identical for ketamine and CGS 19755. Dominant frequencies are computed in 5 minute sliding bins. **A**. A 10 mg/kg dose of ketamine drives oscillations at just over 0.25 Hz for about 1 hour. The dominant frequency has a slight downward trend over this period. **B**. A 12 mg/kg dose of CGS 19755 drives oscillations at 0.4 to 0.5 Hz, starting just over an hour after injection and continuing for at least 4 hours. There is a downward trend in frequency over the first hour.

We next asked how the state of consciousness relates to electrophysiological patterns. Akeju et al. (2016) and Rosen and Hagerdal (1976) have reported that gamma-burst activity appears as patients transition into a behaviorally-defined general-anesthetic state under ketamine ^2,3^. Prior to receiving ketamine, animals left undisturbed and head-fixed in a dark room are typically drowsy, fully closing their eyes from 1/3 to 2/3 of the time, and making saccades and blinks at a rate of 0.2 - 0.4 / sec when their eyes are not fully shut (**Sup. Figs 11-12**). Within minutes of receiving 10 mg/kg of ketamine, animals become lightly anesthetized and are unresponsive to all but very intense stimuli for 40-50 minutes ^12,13^. During this period we observed (in a limited number of sessions for which eye tracking data were collected) that the eyes were partially closed and gaze was fixed in place, with blinks occurring periodically. These blinks were synchronized to the electrophysiological gamma-burst pattern (as seen in **Fig. 1B**, bottom row). Following this period, the eyes opened nearly 100% of the time, and in place of saccades there were brief excursions from center fixation immediately followed by a rapid damped return to central gaze, as if the animals were attempting to saccade but the oculomotor integrator was entirely off-line. This suggests that while transient bursts of activity may reach the oculomotor muscles, they are not followed by any change in the tonic firing rate, that is, it suggests a complete pulse-step mismatch (**Sup. Fig 11E**). Mettens et al. (1990) report the same effect of ketamine in cats. Similar though milder effects were seen after 3 mg/kg (**Sup. Figs. 1, 13**) ^14^. The overall behavioral effect was quite obvious. Even an untrained observer can rapidly identify an animal given either dose, and animals trained in oculomotor tasks fail to initiate even a single trial.

Despite eliciting a similar electrophysiological response, CGS 19755 produces very different behavior. Animals appear bright, alert and responsive to stimuli following 3 and 6 mg/kg doses. Even observers who work with the animals every day were unable to differentiate between animals given 6 mg/kg of CGS 19755 versus a control injection (saline). Several hours after 12 mg/kg doses, animals were slightly slower than usual climbing into their home cage. They also ignored fruit, or put it in their mouths and did not eat it. Normal movement and appetite returned within 12 hours. Like ketamine, CGS 19755 made animals less likely to shut their eyes and increased their blink rate, though these effects were delayed by ∼2 hours (**Sup. Fig. 14**). Unlike ketamine, CGS 19755 had variable effects on saccade frequency.

Animals who received 6 mg/kg of CGS 19755 were able to perform a challenging delayed non-match to sample visually guided saccade task nearly as well as after control injections (**Fig. 4, Sup. Fig. 10**). Meanwhile, animals who received a dose of ketamine with far subtler neural effects (3 mg/kg) did not complete a single trial of the task, and in fact failed to even perform the initial fixation required to initiate a trial.

**Figure 4.**
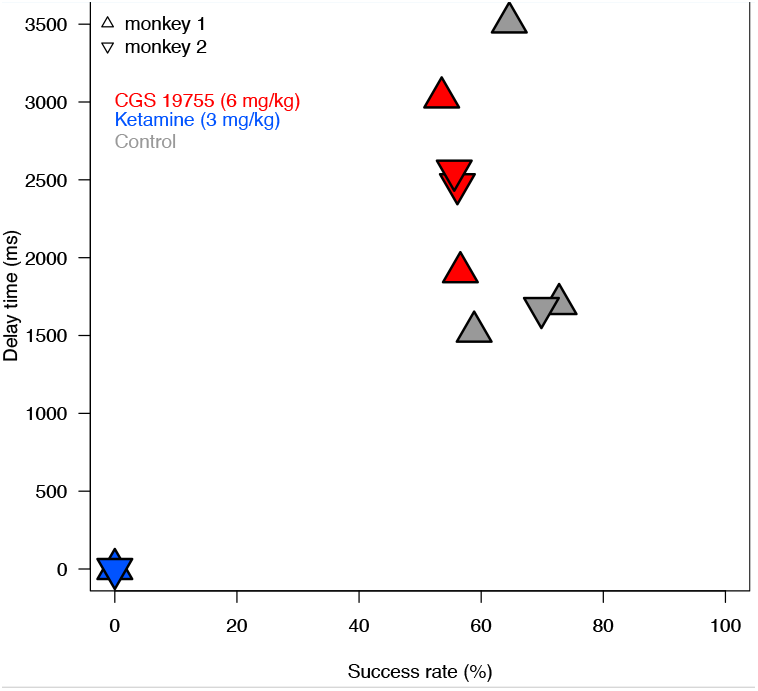
Despite similar electrophysiological signatures in cortex, ketamine and CGS 19755 have very different effects on behavior. Two animals were trained on a difficult 2-target spatial memory task. After a delay, the two targets reappeared along with a novel target, and animals were rewarded for making a saccade to the novel target. The delay was adjusted to maintain behavior at 60-70% correct, typically from 1-3 s. Fifteen minutes after receiving 3 mg/kg ketamine, animals would not perform at any delay length; in fact, they would not even perform a simple fixation task (one session in each of two animals). Two hours after receiving 6 mg of CGS 19755, animals performed at 60% correct with delays of 2-3 s (2 sessions each in each of two animals). Two control trials in each of two animals are shown for comparison. All trials were interleaved, and the operator was blinded to treatment.

## Discussion

We compared the effects of systemic injections of ketamine and the highly selective NMDA receptor antagonist CGS 19755 on intracortical electrophysiological activity and behavior in rhesus macaques. While many of the cortical activity changes produced by anesthetic doses of ketamine (10 mg/kg) also occurred under a sub-anesthetic dose of CGS 19755 (12 mg/kg)—notably including the appearance of gamma bursts—the two drugs differed in their modulation of delta-band power, which was robust under ketamine yet absent under CGS 19755. This difference demonstrates that ketamine’s ability to drive strong delta-band oscillations relies on additional mechanisms beyond NMDA receptor antagonism alone, and points to a key role for the activity underlying delta-band oscillations in causing anesthesia.

The behavioral and electrophysiological effects of ketamine anesthesia we report are in line with previous findings. Ketamine’s behavioral effects are well established, as intramuscular (IM) injections are commonly used to immobilize monkeys for short veterinary procedures and to induce anesthesia prior to the application of an inhaled anesthetic ^12,13^. A 10 mg/kg dose reliably leads to 40 to 50 minutes of light anesthesia, during which time animals are unresponsive to all but the most intense stimuli ^12^. Our eye tracking data show a complete elimination of saccades lasting 40 to 50 minutes following injection (**Sup. Fig. 11, 12**), indicating that animals were successfully anesthetized by the 10 mg/kg ketamine injections used in our experiments.

Cortical activity changes dramatically with induction of ketamine general anesthesia. Within five to ten minutes of injection, multiunit spiking activity, low-pass filtered (0.1 to 1 Hz) local field potential (LFP) voltage, and the low-pass filtered envelope of LFP band-limited power (BLP) all exhibit a repeating two-phase cycle of modulation that is synchronized across all sites recorded in this study (**Figs. 1A-B, 2A-B**). During the “high spiking” phase, spiking activity and the envelope of gamma BLP (24 to 60 Hz) are high while low-pass filtered LFP and the envelope of delta BLP (1 to 4 Hz) are low. During the “low spiking” phase the reverse is true, with little to no spiking activity and a low envelope of gamma BLP, while low-pass filtered LFP and the envelope of delta BLP are high. Each phase of the cycle lasts ∼2s, giving the full cycle of modulation a period of ∼0.25 Hz. These findings align with previous studies of ketamine anesthesia in humans ^2,3^, monkeys ^15^, rodents ^16^, and cats ^15,17-19^. In particular, Akeju et al. (2016) demonstrate that ketamine anesthesia in humans induces “gamma-burst” EEG activity, a cycle of alternating slow-delta (∼0.1–4 Hz) and gamma (∼27–40 Hz) oscillations with a period between 0.1 and 0.2 Hz (Fig. 6 of Akeju et al. 2016). The gamma-burst EEG pattern is specifically associated with a behavioral state of unconsciousness under ketamine anesthesia (lack of response to commands or painful stimuli), while sub-anesthetic ketamine doses cause sustained gamma oscillation uninterrupted by slow-delta oscillations ^3^.

The biological processes underlying ketamine-induced gamma-burst activity and their relationship to the loss of consciousness remain unclear. One possibility is that gamma-burst activity is the result of NMDA receptor antagonism. Homayoun et al. (2007) demonstrate that, in cortex, ketamine preferentially inhibits NMDA receptor activity in GABAergic interneurons, suppressing local negative feedback loops ^20^. This has been proposed as an explanation for the increase in gamma BLP and spiking activity under sub-anesthetic doses of ketamine, and during the high-gamma phase of gamma-bursts under ketamine anesthesia ^3,20,21^. Akeju et al. (2016) hypothesize that the slow-delta (0.1 to 4 Hz) oscillations arise due to a reduction of thalamocortical excitatory input to cortex caused by a blockade of NMDA-mediated excitation of the thalamus by the brainstem. Taken together with the alignment between gamma-burst onset and loss of consciousness, this NMDA-dependent explanation of gamma-bursts is surprising. Ketamine’s dissociative and anesthetic effects are mediated by a variety of molecular targets. These include, but are not limited to, opioid receptors ^22^, monoamine transporters ^23^, hyperpolarization-activated cyclic-nucleotide-gated potassium channel 1 pacemakers ^6^, and muscarinic acetylcholine receptors ^24^. If NMDA antagonism mediates gamma-burst activity, while ketamine general anesthesia depends on activity at one or more of targets besides NMDA, then the close association between gamma-bursts and the loss of consciousness under ketamine may either be coincidental or indicative of an interaction between NMDA antagonism and ketamine’s other targets. Moreover, it should be possible to observe gamma bursts in a conscious animal using only a selective NMDA antagonist.

We show that a 12 mg/kg dose of the selective NMDA receptor antagonist CGS 19755 causes a gamma-burst pattern of activity that is very similar to that seen under ketamine anesthesia. However, CGS 19755 does not produce anesthesia at this dose. The gamma-bursts caused by CGS 19755 occur at ∼0.4 Hz (**Fig. 3**) (compared to ∼0.25 Hz under ketamine, **Fig. 2**). In addition to the high-spikes/high-gamma phase there is also a low-spikes/low-gamma phase during which voltage in the slow LFP range (.1 to 1 Hz) is increased (**Figs. 1C-D, 2C-D**). Critically, the 12 mg/kg dose of CGS 19755 results in an “interruption” of gamma oscillations and spiking activity by increased low-frequency LFP (0.1 to 1 Hz), but not delta BLP (1 to 4 Hz). This raises the possibility that modulation of 0.1 to 1 Hz LFP and 1 to 4 Hz BLP during NMDA-antagonist-induced gamma-burst activity have separate origins. It also suggests that, while interruption of gamma oscillations and spiking activity by delta-band oscillations may reflect an effect of ketamine sufficient to cause anesthesia, interruption by activity in the 0.1 to 1 Hz range does not. The other clear difference in gamma-burst activity with 12 mg/kg CGS 19755 versus 10 mg/kg ketamine is the time to gamma-burst onset (1-2 hours versus 5-10 minutes, respectively). These differences in the electrophysiological effects of CGS 19755 compared to ketamine may either cause, be caused by, or be unrelated to ketamine’s anesthetic effects.

While our findings support the view that delta BLP modulation synchronized with the low-spikes phase of gamma-burst activity sets ketamine anesthesia apart from sub-anesthetic ketamine-induced states (Fig. 1A and C, 2A and C, Supp. Figs. 1 and 2), they do not support the view that delta oscillations arise from local circuit dynamics in the context of reduced thalamocortical excitation. During the low-spikes phase, spiking activity is extremely low - in most cases nearly or completely absent (**Fig. 1**). It is unclear what local circuit dynamics could drive delta oscillations in the absence of the spiking activity on which those dynamics are thought to depend. A more likely explanation is that these delta oscillations reflect afferent activity originating outside of cortex, possibly in the thalamus. This is consistent with the report by Miyasaka et al. (1968) of ketamine-induced thalamic delta oscillations in decorticated cats ^17^. However, the thalamic delta oscillations they observed in decorticated preparations did not have the clear burst-forms observed in their intact brain preparations (periods of high power, similar to what is seen in the monkey cortex during the low-spikes phases in **Fig. 1**). Together, these findings suggest that the gamma-burst-related modulation of delta BLP may depend on activity originating from cortex ^17^. But if this is true, it is unclear why we do not see synchronous delta BLP modulation under CGS 19755, which clearly causes low-frequency modulation of cortical spiking and gamma BLP similar to that seen with ketamine anesthesia.

Simultaneous recordings in this study were limited to posterior cortical regions, which constrains our ability to address the possible interactions between activity in these regions and in frontal cortex or subcortical regions including the thalamus. Future studies recording simultaneously from cortex and thalamus may clarify the interaction between slow modulation of spikes, gamma BLP and low-pass LFP in cortex, and modulation of delta BLP in thalamus and cortex. We avoided higher doses of CGS 19755 due to their prolonged effects and potential neurotoxicity in macaques ^25^, which prevents us from directly addressing the possibility that the full signature of ketamine anesthesia, as identified by Akeju et al. (2016), may be the result of NMDA receptor antagonism alone ^3^. Future experiments could address this by using high doses of CGS 19755 in a lower animal model, or using another selective NMDA antagonist in primates. Because our injections were systemic, we cannot draw definitive conclusions about the mechanisms underlying changes in gamma BLP and spiking activity under ketamine and CGS 19755, although there is reason to believe they arise from inhibition of NMDA receptors on cortical GABAergic interneurons. This might be more clearly addressed using small intracranial injections at individual recording sites.

Akeju et al. (2016) demonstrate that gamma-burst EEG activity occurs with anesthetic doses of ketamine, but not with sub-anesthetic doses. In this report, we have shown that gamma-burst activity can be explained by NMDA antagonism alone, save for the slow (0.1 to 1 Hz) synchronized modulation of delta BLP. Critically, we also show that prolonged NMDA-antagonist-induced gamma-burst activity can occur without anesthesia, and is therefore not sufficient to cause the anesthetic effects of ketamine.

## Materials & Methods

### Animals and behavior

Three macaques served as subjects in this study (here labeled monkeys E, G and I). Animals were cared for and handled in accordance with the Guide for the Care and Use of Laboratory Animals, and all procedures were approved by the Washington University IACUC. During recording, macaques were fully hydrated and sat head-fixed in a dark room. Behavior was unconstrained, and the animals had no expectation of a task or reward. Halfway through each recording, animals were given a systemic injection of ketamine (1, 3 or 10 mg/kg) or CGS 19755 (3, 6 or 12 mg/kg).

### Recording

A total of 29 sessions were recorded with an average duration of 4.6 hours: 7 from monkey E, 17 from monkey G, and 5 from monkey I. Intracranial electrophysiological signals were recorded simultaneously from three to eight locations in each hemisphere. Recordings were collected from the Lateral Intraparietal area (LIP), the Medial Superior Temporal area (MST), and areas 31, 23B, V4, 7A, V3D, and V3A. Results were generally similar for the three monkeys, and thus the data were combined.

Electrodes were targeted to each area of interest using anatomical MRI images ^26^. Briefly, each animal’s brain was accessed via bilateral 15 mm (internal diameter) chronic custom recording chambers. T1 weighted MRI images (MPRAGE; .5 mm isotropic voxels) were obtained using a custom phantom in the chamber that provides visualization of the chamber and allows for the virtual projection of a chamber-based coordinate system down into the brain. A 3D atlas of the macaque brain based on the Saleem and Logothetis D99 macaque atlas was nonlinearly aligned to each animal’s T1 image, allowing us to map positions in the chamber-based coordinate system to atlas-defined anatomical regions and vice versa ^27-30^. In one monkey, a postmortem T1 was collected with recording electrodes still in the brain to validate targeting accuracy by comparing their expected locations to their actual locations in the MRI image.

Extracellular voltage signals (local field potential, LFP) were collected using tungsten microelectrodes (Alpha Omega LTD) attached to motorized drives (NAN). LFP signals were recorded by a digitizing amplifier (Intan), and an Open Ephys acquisition system ^31^. The amplifier sampled each signal at 30 kHz, and was configured with a 0.02 Hz 1-pole high-pass analog filter. In a limited number of sessions, pupil position (horizontal and vertical) and pupil diameter (vertical) were collected using an infrared camera system (EyeLink), and recorded as three separate analog inputs through the same Open Ephys acquisition board used to collect LFP data.

### Behavioral Task

Two additional monkeys (H and F) were asked to perform a challenging 2-target spatial memory task under the influence of Ketamine, CGS-19755, or a control injection of saline. In this task, two targets were presented, disappearing after a certain period and re-emerging along with a novel target. The animals were rewarded when they made a saccade towards the novel target. The delay period between target disappearance and reappearance was carefully adjusted to maintain the monkeys’ performance within the 60-70% accuracy range, typically extending from 1.5 to 3.5 seconds. For all behavioral sessions, animals were given one injection at two hours and one injection fifteen minutes before being asked to engage in the task. In non-control sessions, only one of the injections would contain a drug. In total, ten sessions were conducted: six sessions with monkey H and four sessions with monkey F. Among these, four sessions (three for monkey H and one for monkey F) were control conditions. Two sessions for each monkey were conducted under the influence of CGS 19755 (6 mg/kg at -2 hours) and one session for each monkey was performed under the effect of ketamine (3 mg/kg at -15 minutes). All trials were interleaved, and the person administering the injections was blind to the treatments.

### Analysis

Analyses were performed with custom software written in Julia ^32^, Python, and Matlab (MathWorks). Electrophysiological signals were processed separately offline to extract single- and multi-unit spiking activity. First, each signal was bandpass filtered between 500-3000 Hz, and points where the filtered signals value crossed a threshold of ±4 standard deviations were set aside as potential spikes. These were divided into groups using the “masked EM” clustering algorithm implemented in KlustaKwik ^33^, and these groups were manually reviewed to reject artifacts and clusters that appeared or disappeared at any point during the recording window, especially around the time of drug injections. Spike times were converted into spike rate estimates by convolving their time-binned representation with a gaussian kernel (s.d. = 84ms, doubling or halving kernel width does not effect outcomes of downstream analyses). For all other analyses, LFP signals were first decimated to 250 Hz and notch filtered at 60 Hz to remove power line noise.

The spectrograms in **Fig. 1A,C** were calculated using the filter-Hilbert method. All filters were noncausal 5-pole Chebyshev Type-II filters with 20 dB stopband ripple. A total of 40 filters were used, with center frequencies logarithmically spaced from 2-100 Hz (inclusive) and bandwidth for each filter set at 30% of its center frequency. To emphasize low-frequency changes in power, the power timecourse for each band was then band-pass filtered from 0.1-1 Hz with a 9-pole Chebyshev Type-II filter with 40 dB stopband ripple, and normalized via Z-transform using its pre-injection mean and standard deviation. For analyses using canonical frequency bands, LFP was band-pass filtered to obtain gamma (30-60 Hz), delta (1.5-4 Hz) and “slow” (0.1-1 Hz) LFP using 9-pole Chebyshev Type-II filters with 40 dB stopband ripple. Gamma and delta band-limited power (BLP) timecourses were obtained by applying the Hilbert transform to gamma and delta LFP, then band-pass filtered from 0.1-1 Hz to emphasize low-frequency changes in power as described above. For **Fig 1B,D** gamma BLP and delta BLP were Z-transformed using pre-injection mean and standard deviation.

The intensity of low-frequency signal modulation (**Fig. 2A,C**) was quantified by taking the power of each signal between 0.1 and 1 Hz. Signals (spike rate, slow LFP, gamma BLP and delta BLP) were decimated to 4 Hz then filtered using a 9-pole Chebyshev Type-II filter with 40 dB stopband ripple, followed by the Hilbert transform. The resulting low-frequency power time series were normalized with a Z-transform using pre-injection mean and standard deviation, aligned by injection time, and averaged across subjects and sessions separately for each drug condition.

The windowed intra-modal coherence between simultaneously recorded low-pass LFP and low-passed (0.1-1 Hz) BLP envelope (**Fig. 2B,D**) was obtained by decimating each signal to 4 Hz, applying a 0.1-1 Hz filter (in the case of the BLP signals but not the already filtered low-pass LFP signal), then calculating coherence in 60-second non-overlapping windows aligned to the time of injection (mt_coherence() from DSP.jl) ^34^. The Fisher z-transformation was applied to the resulting time-frequency data, which were then normalized by subtracting the pre-injection mean and dividing by the pre-injection standard deviation separately for each frequency. Data were then averaged across subjects and sessions separately for each drug condition.

To calculate the dominant frequency of modulation over time (**Fig. 3**), we first calculated the autocorrelation of each signal within 5-minute windows beginning after injection, then took the reciprocal of the mean period of the tallest peaks for each window to obtain a dominant frequency for each signal for each window. Data were then averaged across subjects and sessions separately for each drug condition, and timepoints before and after the drug’s peak effect were manually excluded for display purposes.

## Data Availability

The data that support the findings of this study are available from the corresponding author upon reasonable request.

## Code Availability

The code that analyzed the neural data of this study are available from the corresponding author upon reasonable request.

## Acknowledgements

This work was supported by the National Institute of Mental Health at the National Institutes of Health (grant numbers R34NS118618, R01MH128286); the National Institute of General Medical Sciences at the National Institutes of Health (grant number T32GM008151); the National Eye Institute at the National Institutes of Health (grant number R01EY012135) and the McDonnell Center for Cellular and Molecular Neurobiology (grant number 3930-26275F).

## Author Information

**Affiliations**

*Department of Neuroscience, Washington University School of Medicine, St Louis, MO, 63110, USA*

Benjamin T. Acland, Lawrence H. Snyder

*Department of Anesthesiology, Washington University School of Medicine, St Louis, MO, 63110, USA*.

Ben Julian A. Palanca

*Department of Psychiatry, Washington University School of Medicine, St Louis, MO, 63110, USA*.

Ben Julian A. Palanca

*Department of Radiology, Washington University School of Medicine, St Louis, MO, 63110, United States*

Janine Bijsterbosch

*Department of Biomedical Engineering, Washington University, St Louis, MO, 63130, USA*

Lawrence H. Snyder

**Contributions**

B.T.A. and L.H.S. designed research and analyzed data; B.T.A. performed research; and B.T.A., B.J.A.P., J.B. and L.H.S. wrote the paper.

## Competing Interests

The authors declare no competing interests.

**Corresponding Author**

Correspondence to Lawrence H. Snyder

