## Supplemental Figures for "Gamma-burst cortical activity in awake behaving macaques"

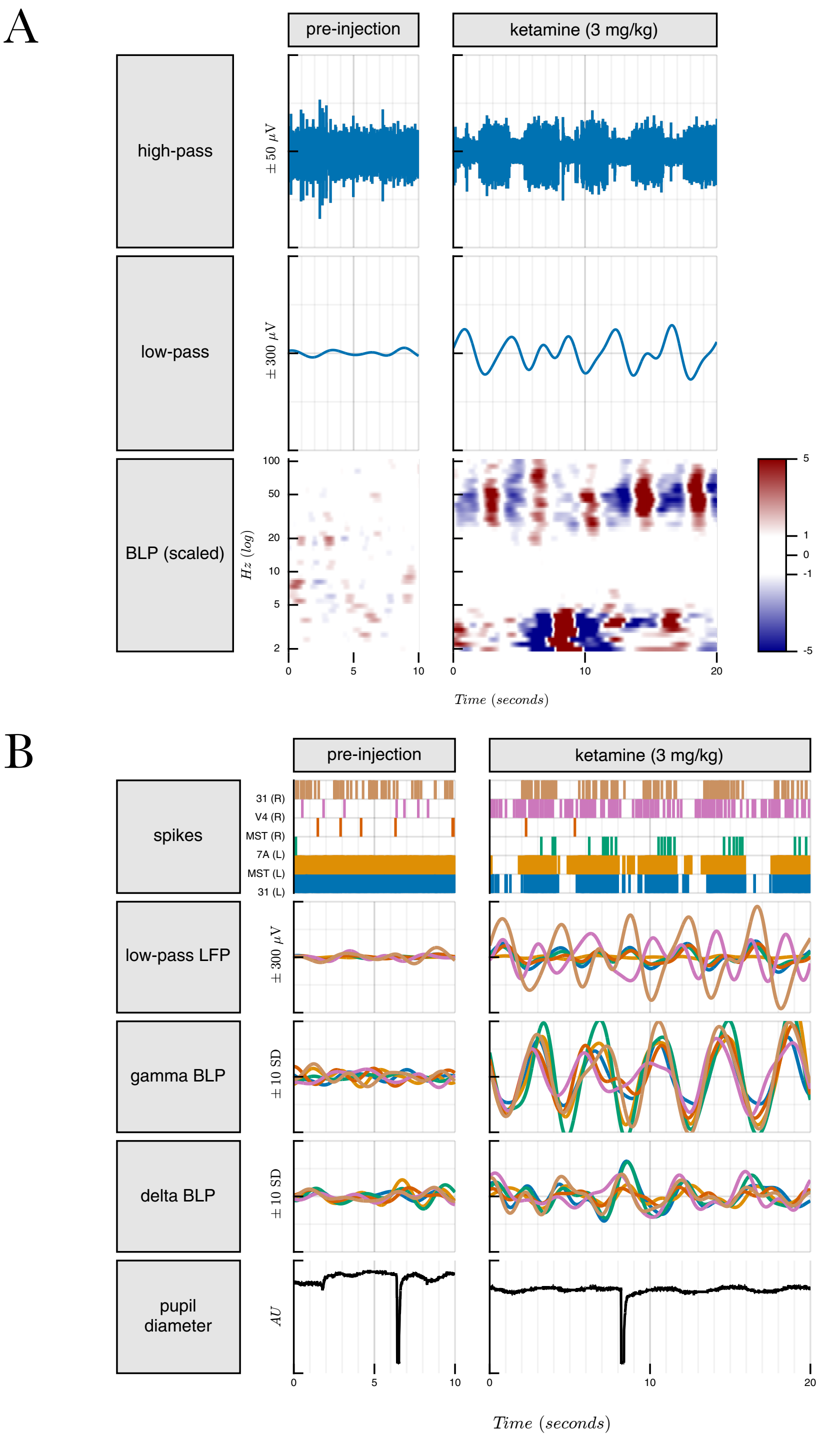


**Supp. Figure 1.** Similar to Figure 1, panels A and B, with 3 mg/kg ketamine.

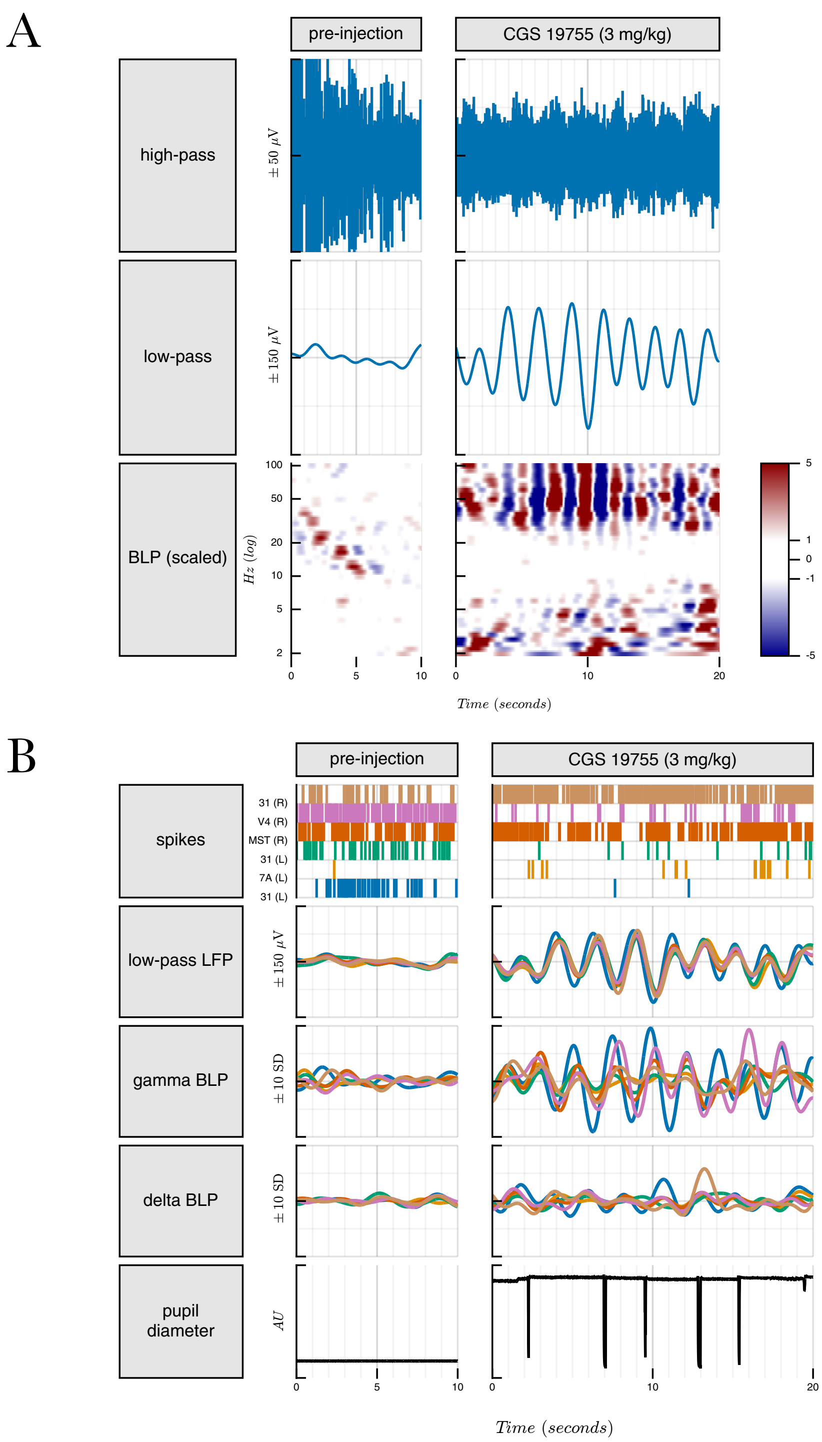


**Supp. Figure 2.** Similar to Figure 1, panels C and D, with 3 mg/kg CGS 19755.

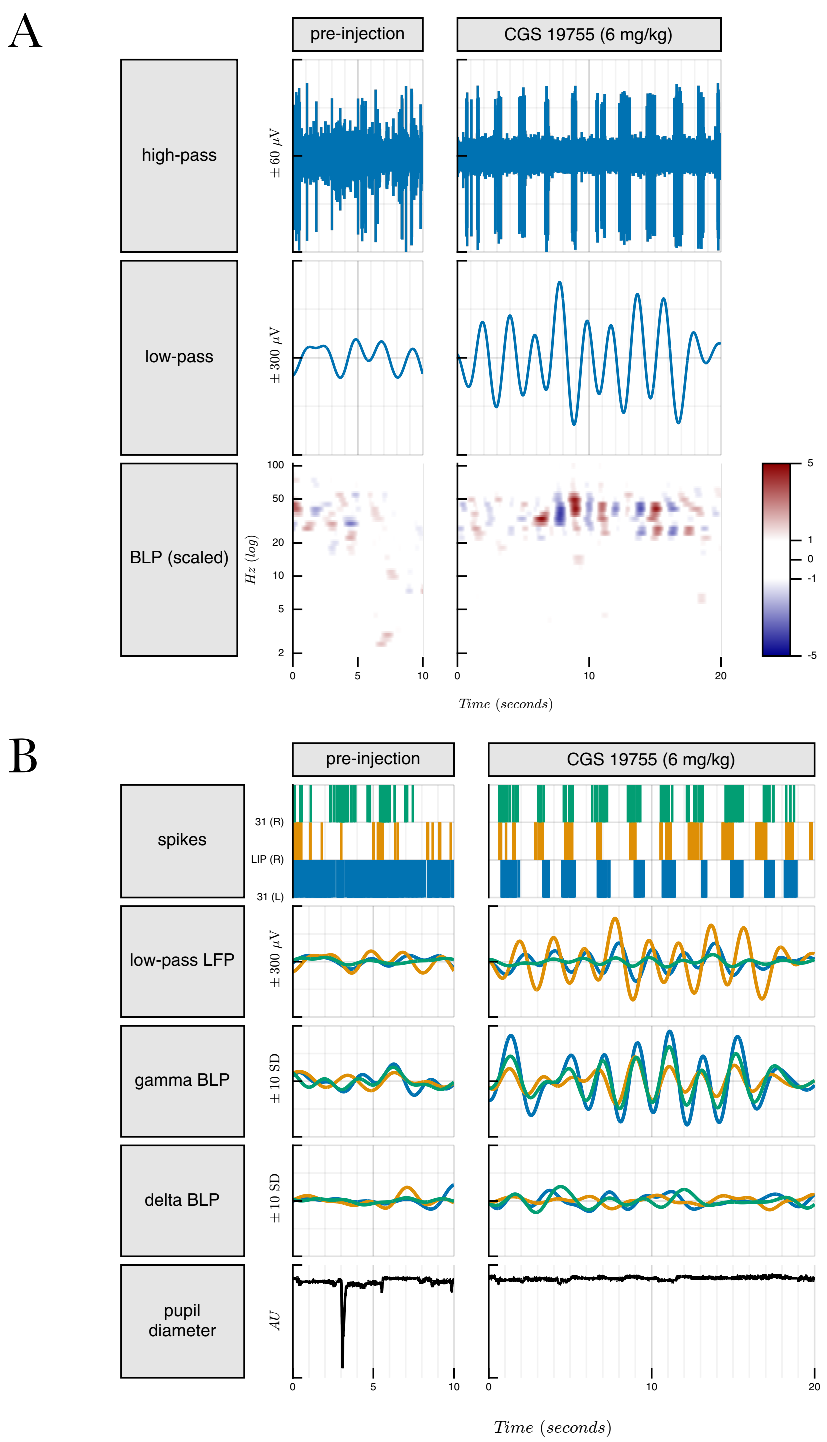


**Supp. Figure 3.** Similar to Figure 1, panels C and D with 6 mg/kg CGS 19755.

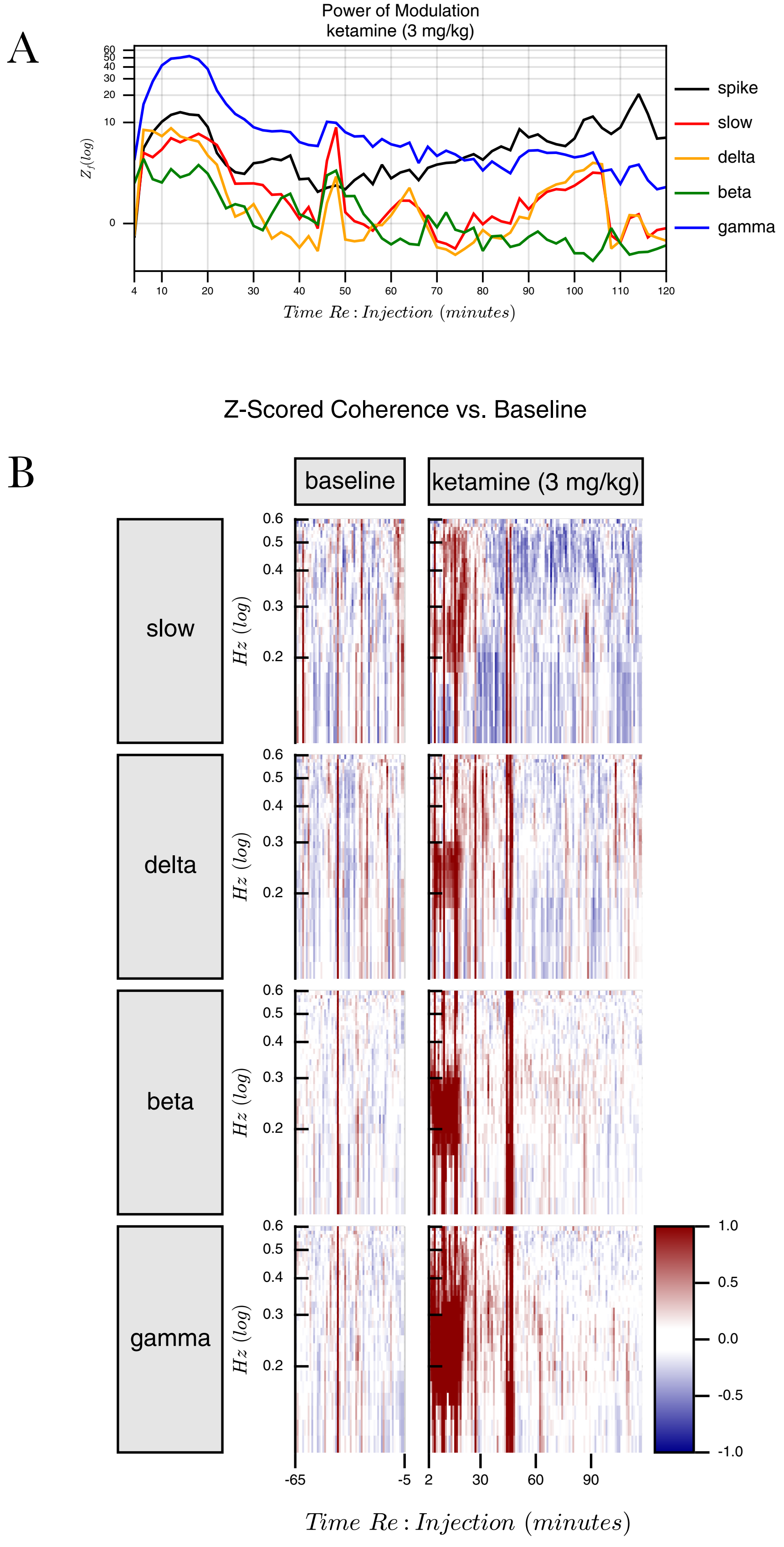


**Supp. Figure 4.** Similar to Figure 2, panels A and B, with 3 mg/kg ketamine.

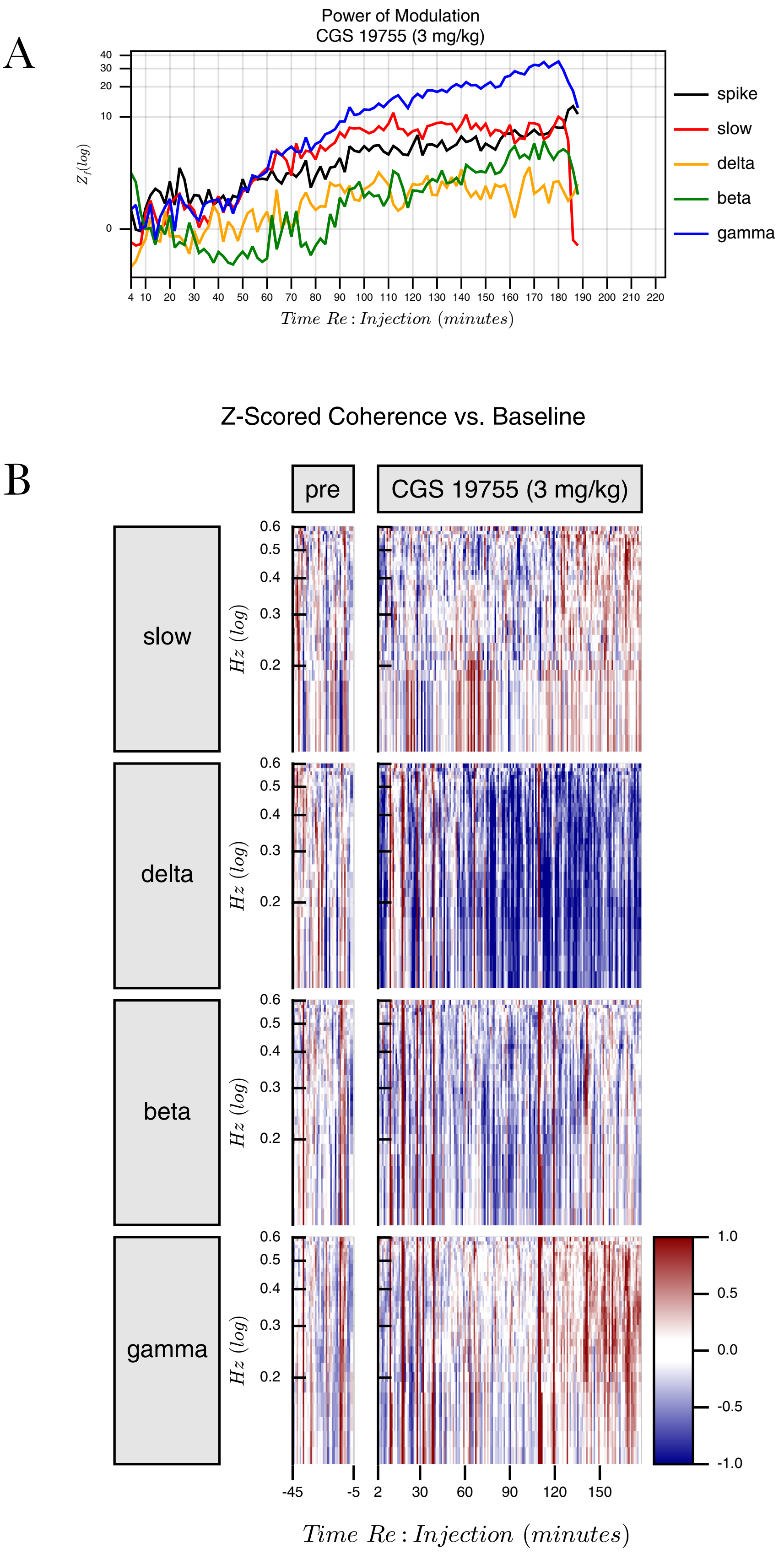


**Supp. Figure 5.** Similar to Figure 2, panels A and B, with 3.2 mg/kg CGS 19755.

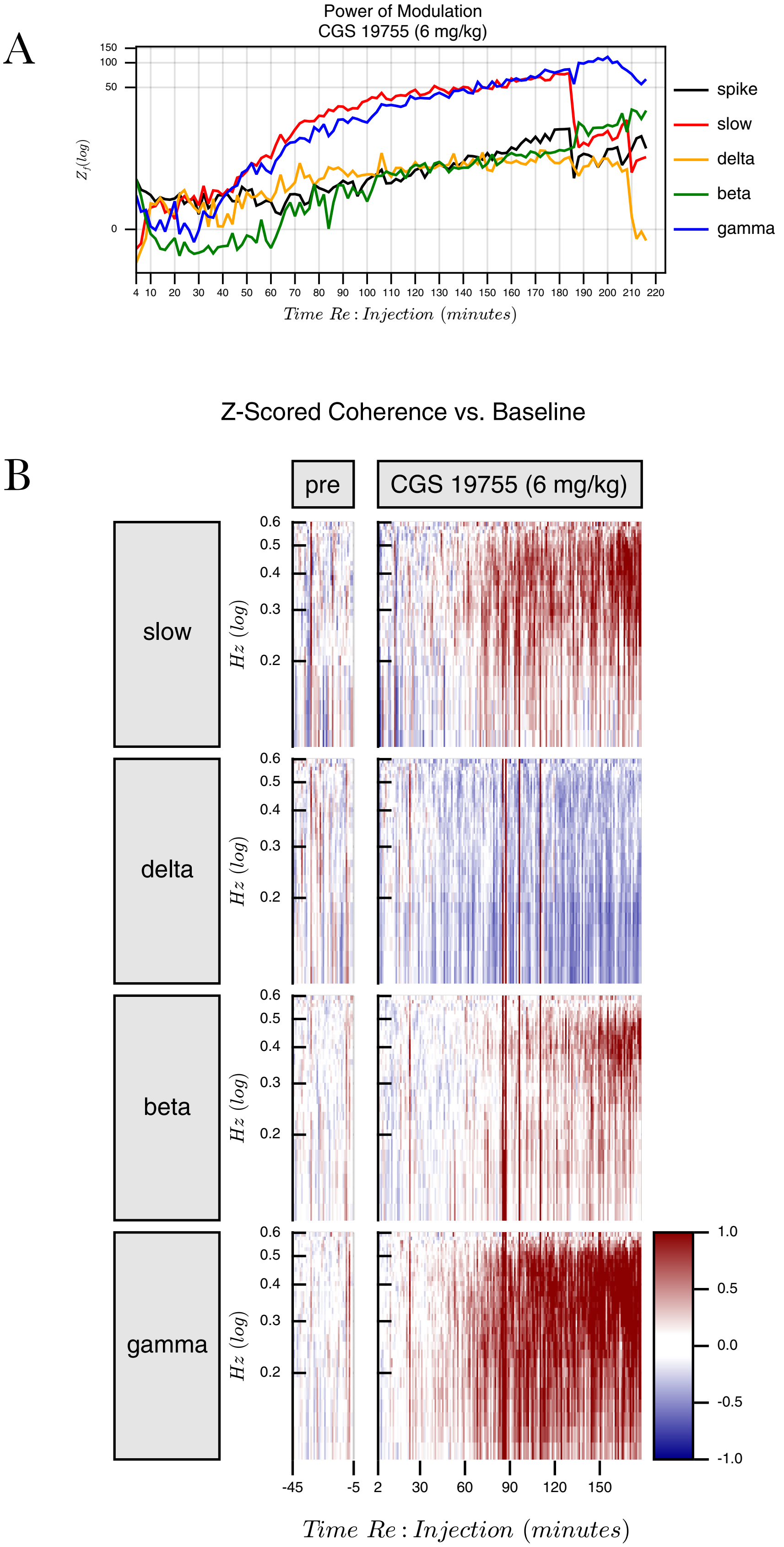


**Supp. Figure 6.** Similar to Figure 2, panels A and B, with 6 mg/kg CGS 19755.

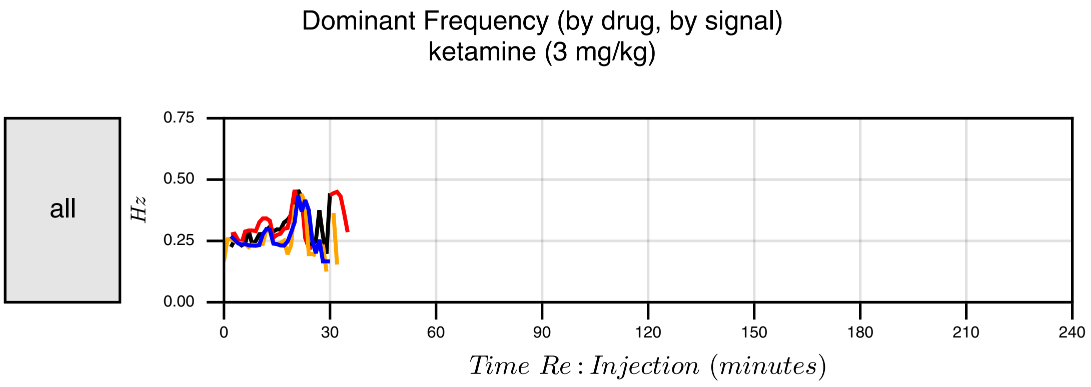


**Supp. Figure 7.** Similar to Figure 3, panel A, with 3 mg/kg ketamine.

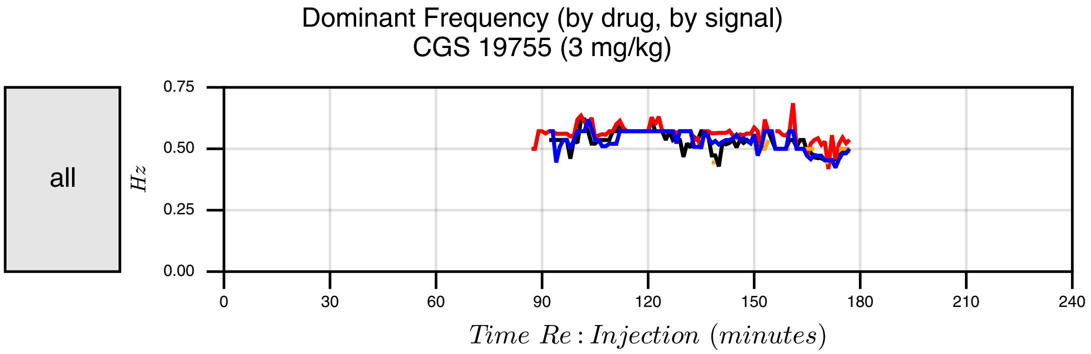


**Supp. Figure 8.** Similar to Figure 3, with 3.2 mg/kg CGS 19755.

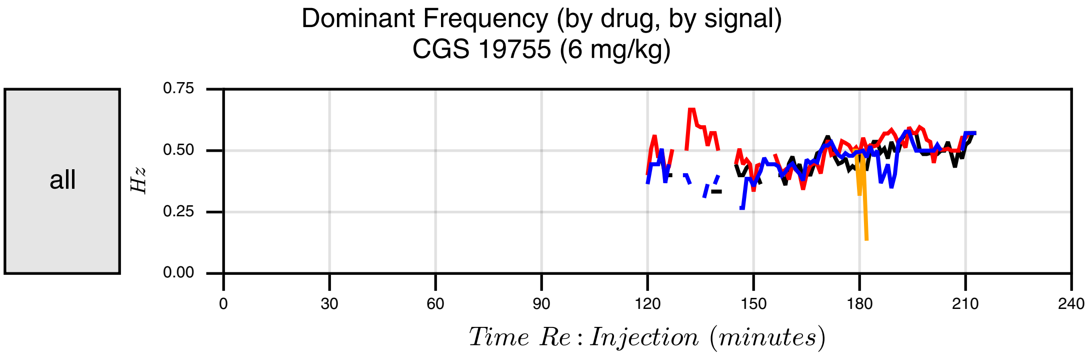


**Supp. Figure 9.** Similar to Figure 3, with 6 mg/kg CGS 19755.

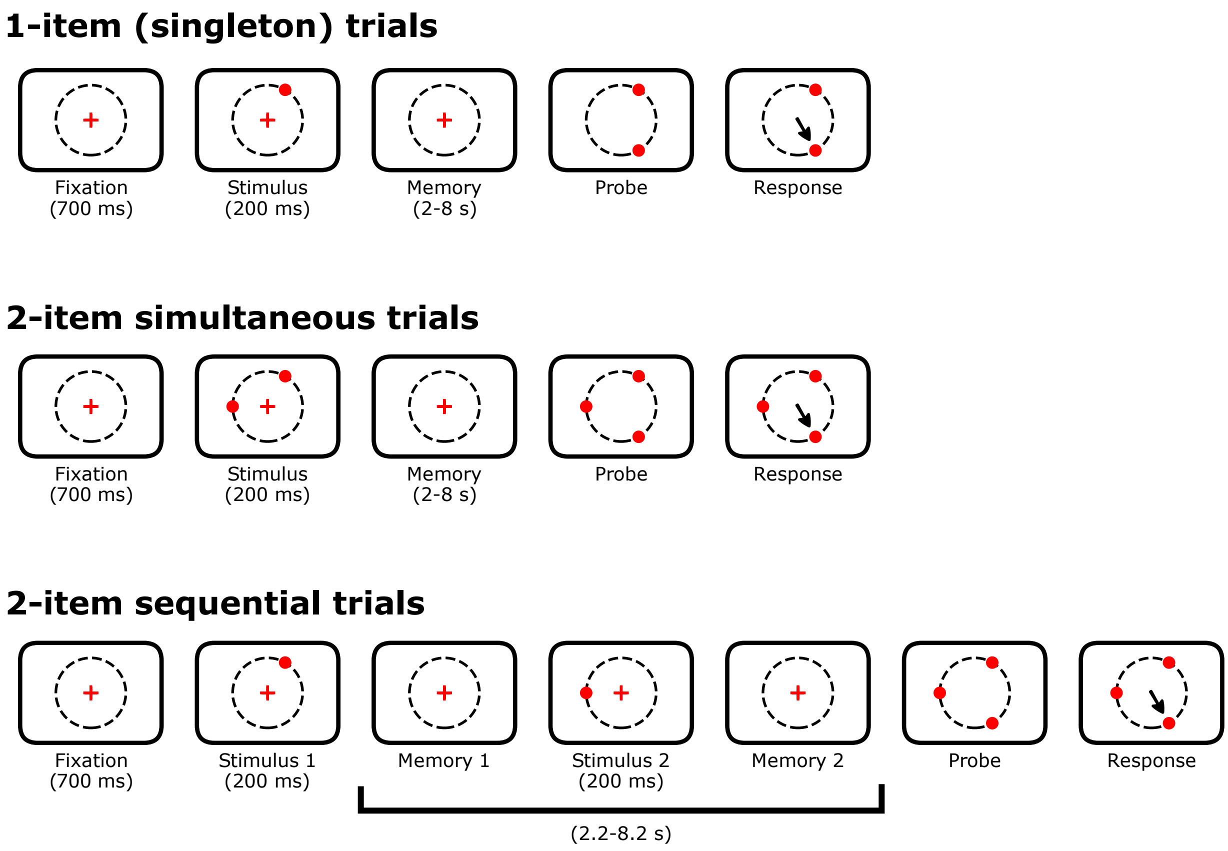


**Supp. Figure 10.** The spatial memory tasks used in association with Figure 4. Data in that figure are for “2-item sequential” trials; results for the other two trial types are similar.

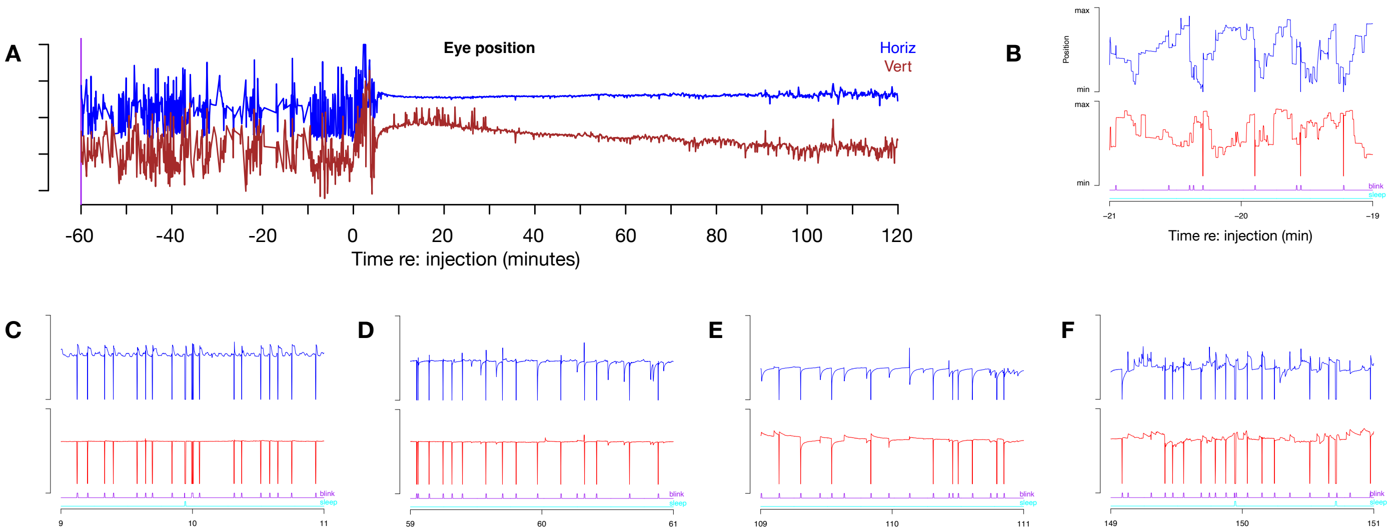


**Supp. Figure 11. A.** Time course of eye position before and after a 10 mg/kg ketamine injection. Vertical scale is arbitrary. Because the time scale is so compressed, the eye position traces when the eyes are moving are distorted (e.g., aliased). Panels B-F each show 2 minute intervals of data to overcome this. **B.** Baseline: clear saccades, no drift, 4.5 blinks/minute. (Blinks often result in artifactual downward eye position deflections.) Vertical scale is arbitrary, but constant for B-F. **C.** Acute: eyes pinned at center. No drift, no saccades. (Vertical wobbles are cylical lid droop, synchronized to the 0.25 Hz spike and LFP cycles). 9 blinks/min **D.** 1 hour: eyes mostly pinned at center, but ~7 downward shifts followed by rapid gaze paretic drift back to center. 8 blinks/min **E.** 2 hours: eyes close to center, but with multiple rapid deflections (saccades) followed by exponential drift back to center (gaze paretic nystagmus). 10 blinks/min, 2 short sleep episodes. **F.** 1.5 hours: eyes still mostly pinned at center, but with multiple downward, left and right movements followed by exponential drift back to center. 8 blinks/min.

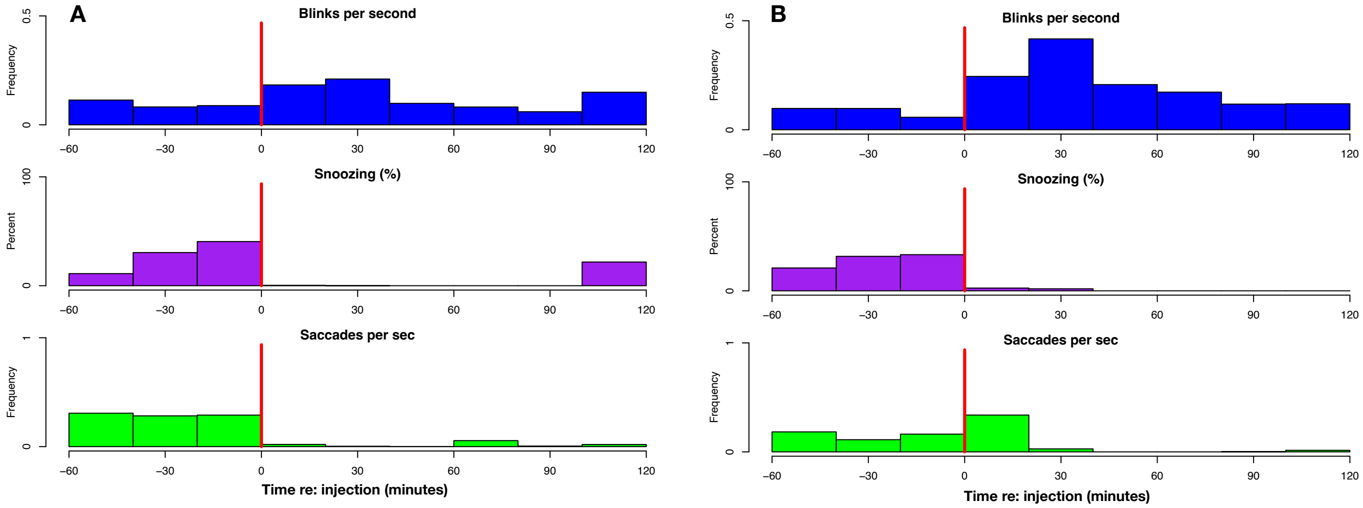


**Supp. Figure 12.** Two 10 mg/kg ketamine sessions from two different animals showing an increase in blink rate, a drop in periods in which the eyes remained closed for longer than a slow blink ("snoozing", eyes shut for >250 ms), and a drop in the frequency of saccades. (In session B saccade frequency increase for ~5 minutes after injection and then dropped sharply— see previous supplemental figure, panel A).

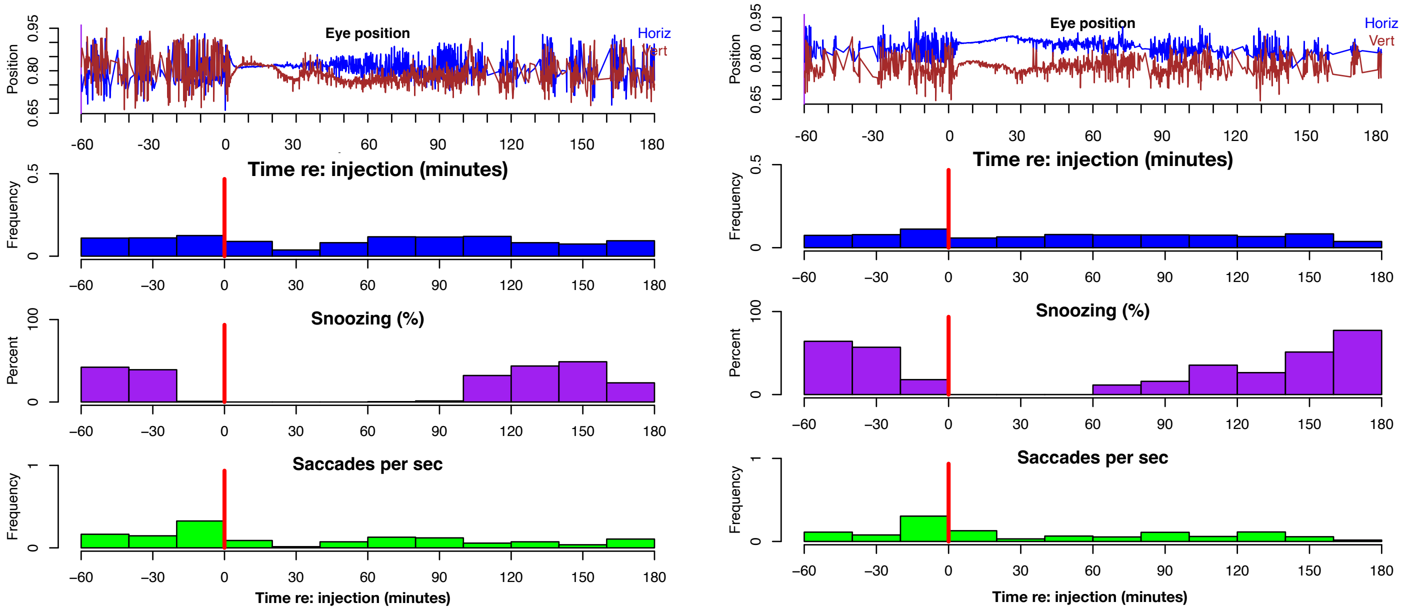


**Supp. Figure 13.** Two 3 mg/kg ketamine sessions from two animals showing a drop in periods in which the eyes remained closed for longer than a slow blink ("snoozing", eyes shut for >250 ms) and a drop in the frequency of saccades. There was minimal effect on blink rates, and behavior began to recover more quickly than after 10 mg/kg injections (compare with previous figure).

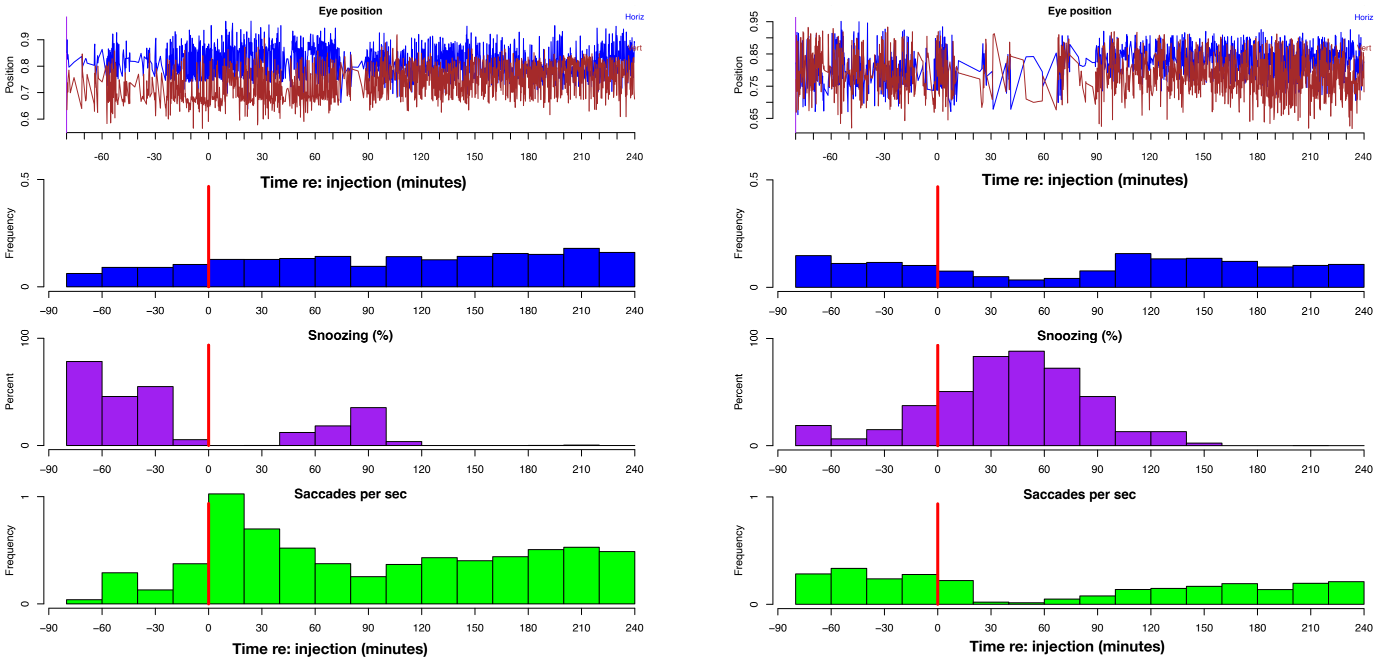


**Supp. Figure 14.** CGS 19755 sessions (12 mg/kg) from two animals. Both animals stop “snoozing” (eyes closed for longer than the duration of a blink) starting 2 hours after the injection. In the late period, saccade and blink frequency increase in one animal, and in the other, blink rate stays constant while saccade rate drops slightly.
